# Classification of grain amaranths using chromosome-level genome assembly of ramdana, *A. hypochondriacus*

**DOI:** 10.1101/2020.06.27.174920

**Authors:** Saptarathi Deb, Suvratha J, Samathmika Ravi, Raksha Rao K, Saurabh Whadgar, Nivedita Hariharan, Shubham Dixit, Meeta Sunil, Bibha Choudhary, Piergiorgio Stevanato, Eswarayya Ramireddy, Subhashini Srinivasan

## Abstract

In the age of genomics-based crop improvement, a high-quality genome of a local landrace adapted to the local environmental conditions is critically important. Grain amaranths produce highly nutritional grains with a multitude of desirable properties including C4 photosynthesis highly sought-after in other crops. For improving the agronomic traits of grain amaranth and for the transfer of desirable traits to dicot crops, a reference genome of a local landrace is necessary. Towards this end, our lab had initiated sequencing the genome of *Amaranthus (A.) hypochondriacus* (A.hyp_K_white) and had reported a draft genome in 2014. We selected this landrace because it is well adapted for cultivation in India during the last century and is currently a candidate for TILLING-based crop improvement. More recently, a high-quality chromosome-level assembly of *A. hypochondriacus* (PI558499, Plainsman) was reported. Here, we report a chromosome-level assembly of A.hyp_K_white (AhKP) using low-coverage PacBio reads, contigs from the reported draft genome of A.hyp_K_white, raw HiC data and reference genome of Plainsman. The placement of A.hyp_K_white on the phylogenetic tree of grain amaranths of known accessions clearly suggests that A.hyp_K_white is genetically distal from Plainsman and is most closely related to the accession PI619259 from Nepal (Ramdana). Furthermore, the classification of another accession, Suvarna, adapted to the local environment and selected for yield and other desirable traits, is clearly *A. cruentus*. A classification based on hundreds of thousands of SNPs validated taxonomy-based classification for a majority of the accessions providing the opportunity for reclassification of a few.

## INTRODUCTION

Grain amaranth, also known as Ramdana (The God’s grain) or Rajgira or Rajeera, has been in continuous cultivation at least since last century in India. This crop was declared “The Future Crop” by the United States in the 1980s based on a decade of intense research in the 1980s (National Research Council, Amaranth: Modern Prospects for an Ancient Crop, National Academy Press, Washington, DC, 1986). At a time when gluten-free, protein-rich, high-fiber and high nutritional values are becoming attractive labels in supermarkets around the globe, grain amaranths deserving all these labels cannot be ignored as a future crop. Furthermore, desirable agronomic traits including drought resistance, C4 photosynthesis, herbicide resistance and high dry-biomass renders grain amaranths as a potential model organism by researchers working on the improvement of other edible dicots. In the context of increasing demand on water and other natural resources from an increasing world population, grain amaranths offer an alternative to other staple cereals such as rice or wheat. With one-sixth of the world population under the poverty line, the value of the seed protein content in amaranth to India cannot be overestimated. Unfortunately, despite India being one of the few countries where multiple landraces of grain amaranths are under continuous cultivation for more than a century, they have received little attention and failed to reach the status of a staple crop.

Interestingly, grain amaranths, domesticated around 8000 years ago, enjoyed equal status as corn during the Aztec and Inca civilizations^1^. This practice went into oblivion after the Columbian exchange. It took about 500 years after the Columbian exchange and intense efforts by the US before this magic grain received the much-deserved global attention. A decade-long research conducted by the Rodale Institute during the 1980s enabled the creation of more than 800 species/varieties, which are currently maintained in a germplasm (GRIN-Global). Interestingly, this germplasm includes seeds from many amaranth landraces from South Asia including India. It is believed that these landraces, which are in contiguous cultivation in distal geographical locations in India, have already adapted to diverse environmental conditions prevalent in Nepal, as well as in East and South India.

More recently, the plummeting cost of sequencing has democratized the application of genomics technologies not only to non-model crops but has extended its reach to individual landraces with direct benefit to local farmers. In this context, the draft assembly of a landrace from India was sequenced and reported^2^. This landrace was selected for its aggressive growth, and its yield compared to a few other landraces including one with red inflorescence cultivated in India. Since then, the chromosome-level genome of a different cultivar with an accession of PI558499 (Plainsman), has been deciphered using state-of-the-art technologies including Bionano, HiC, and long PacBio reads^3^. This high-quality assembly has now allowed placement of about one hundred accessions from the germplasm on a phylogenetic tree^4^ allowing for both establishing genotype-to-phenotype relationships and to place various landraces with very distinct phenotype on the tree for further characterization.

Giving chromosomal context to genes and other genetic elements is one of the most sought-after goals in genome assembly. While the genomes of hundreds of organisms at the draft stage allow deciphering the majority of the proteomes, draft genomes lack chromosomal context under which they evolve and transcribe, which is necessary for a full understanding of biology. Before long-read sequencing became commonplace, experimentally generated mate-pair reads of increasing insert sizes were routinely used to generate scaffolds from contigs. Tools, such as SOAPdenovo, use the known insert size between the mate-pair reads to connect contigs into longer scaffolds by filling the gaps with Ns^5^. Such an approach can simply be extended for reference-guided improvement of draft genomes of a plant using simulated mate pairs of varying insert sizes from an existing assembly of a different variety/cultivar of the same species. For example, mate-pair libraries from one *Arabidopsis thaliana* strain were shared across many strains to build super-scaffolds for all individuals^6^. Also, assisted assembly of closely related species significantly improved the contiguity of low coverage mammalian assemblies^7^. The draft genomes of four species including bushbaby, African elephant, rabbit and guinea pig from the “Mammal24 - 2X” project were built using both human and canine references^7^. More recently, our group demonstrated that two draft genomes of the same species could be used to mutually improve scaffolding of the genome of *Anopheles stephensi* to the point that a set of low-resolution physical markers was sufficient to build the chromosomes^8^.

The utility of mate-pairs from one strain to build the scaffolds for the other require DNA level similarity, which is often not the case even for closely related species. This is because DNA diverges faster even between very closely related species. However, natural selection puts sufficient selection pressure on protein sequences for maintaining functional contiguity required during evolution. In this case, one could use synteny between species at protein levels to build chromosomes. Recently, a chromosome level genome of *Lates calcarifer* was assembled from a draft genome using long-read sequencing, transcriptome data, optical/genetic mapping and synteny to two closely related seabasses^9^. In yet another report, 16 out of 60 chromosomes of the Tibetan antelope were reconstructed from draft assemblies using its homology to cattle^10^. In fact, using independent mapping data and conserved synteny between the cattle and human genomes, 91% of the cattle genome was placed onto 30 chromosomes^11^. In a review article, synteny has been used to filter, organize and process local similarities between genome sequences of related organisms to build a coherent global chromosomal context^12^. Similarly, the malarial strain, *Plasmodium falciparum* HB3, was improved using the reference of *P. falciparum* 3D7 combined with an assisted assembly approach that significantly improved the contiguity of the former^7^.

Grain amaranth is yet to reach an agronomic status in India. While a large number of landraces, adapted to local environments for small scale cultivation exist, their origins and relations to the large germplasm, collection at GRIN-Global are not established. For genomics-based crop improvement of local landraces, it is critical to classify these with respect to accession from the germplasm collection. More recently, using genotyping-by-sequencing (GBS), 94 accessions for grain amaranths have been classified^4^. It is of interest to decorate this phylogenetic tree with landraces of importance to India and elsewhere. While GBS is a cost-effective technology for classifying large number of accessions, it covers only 10% of the genome, which depends on the sample preparation protocol and reagents used. For a small number of landraces it is not trivial to generate the sequences of the same 10% by reproducing the protocol/ reagents, which challenges the placement of additional varieties on the phylogenetic tree generated by GBS. However it is straight-forward to generate whole genome sequencing data, which challenges the placement of additional varieties on the phylogenetic tree. There is a need for normalizing WGS data with GBS data to aid classification of additional landraces from India and elsewhere.

In this paper, we report a de novo assembly of a landrace (A.hyp_K_white) and demonstrate that, in the presence of a reference genome for a distal variety, a chromosome-level assembly can be generated at a reasonable cost. Also, we normalized the variants from GBS and WGS data for various accessions enabling decoration of the phylogenetic tree including many accessions with the landraces of interest from India.

## RESULTS

### Assembly of the landrace A.hyp_K_white

As shown in the flowchart in Figure 6, PacBio reads using RSII technologies sequenced in 2013 with an average length of 7.5 kb with a coverage of 25X for A.hyp_K_white were assembled using state-of-the-art tools CANU^13^ and FLYE^14^ to obtain an assembly with L50 values of 1395 and 944 respectively. These two assemblies were then merged using Quickmerge^15^ to improve the L50 to 623. This was further improved by merging the Illumina assembly from our previously reported draft genome of the same landrace A.hyp_K_white^2^, to get a contig-level assembly with an L50 of 593 (AhK593). We used simulated mate pairs from the reference genome of the Plainsman strain^3^, to build scaffolds of contigs from AhK593 to an L50 of 56 (AhK56) and subsequently using raw HiC data of the Plainsman strain from public sources to obtain a scaffold-level assembly with an L50 of 20 (AhK20) using SALSA^16^. Scaffolds from AhK20 are further stitched based on synteny to A.hyp.V.2.1 to get the final chromosome-level assembly AhKP for the accession A.hyp_K_white. Figure 1.a shows the synteny of the scaffolds from the assembly AhK20 on A.hyp.V.2.1 and Figures 1.b, 1.c and 1.d show synteny of AhKP on to A.hyp.V.2.1 in various representations. Table 1 shows the assembly statistics

**Table 1:** Assembly statistics

| Name | AhK593 | AhK99 | AhK56 | AhK20 | AhKP |
| --- | --- | --- | --- | --- | --- |
| Number of contigs | 4796 | 2960 | 1926 | 1678 | 16 |
| Longest scaffold (Mb) | 1.83 | 7.00 | 11.41 | 24.01 | 39.67 |
| L50 | **593** | **99** | **56** | **20** | **7** |
| N50 (Mb) | 0.19 | 0.89 | 2.00 | 5.40 | 23.02 |
| Assembly size (Mb) | 418.25 | 419.17 | 408.03 | 408.19 | 388.93 |
| Number of ATGCs | 401,504,412<br>(95.99 %) | 401,504,412<br>(95.78 %) | 366,961,434<br>(89.93 %) | 366,961,434<br>(89.89 %) | 348,890,439<br>(89.70 %) |
| Number of Ns | 1,671,100 (4.00 %) | 17,670,600 (4.21 %) | 41,071,762 (10.06 %) | 41,229,762 (10.10 %) | 40,041,419 (10.29 %) |

**Figure 1:**
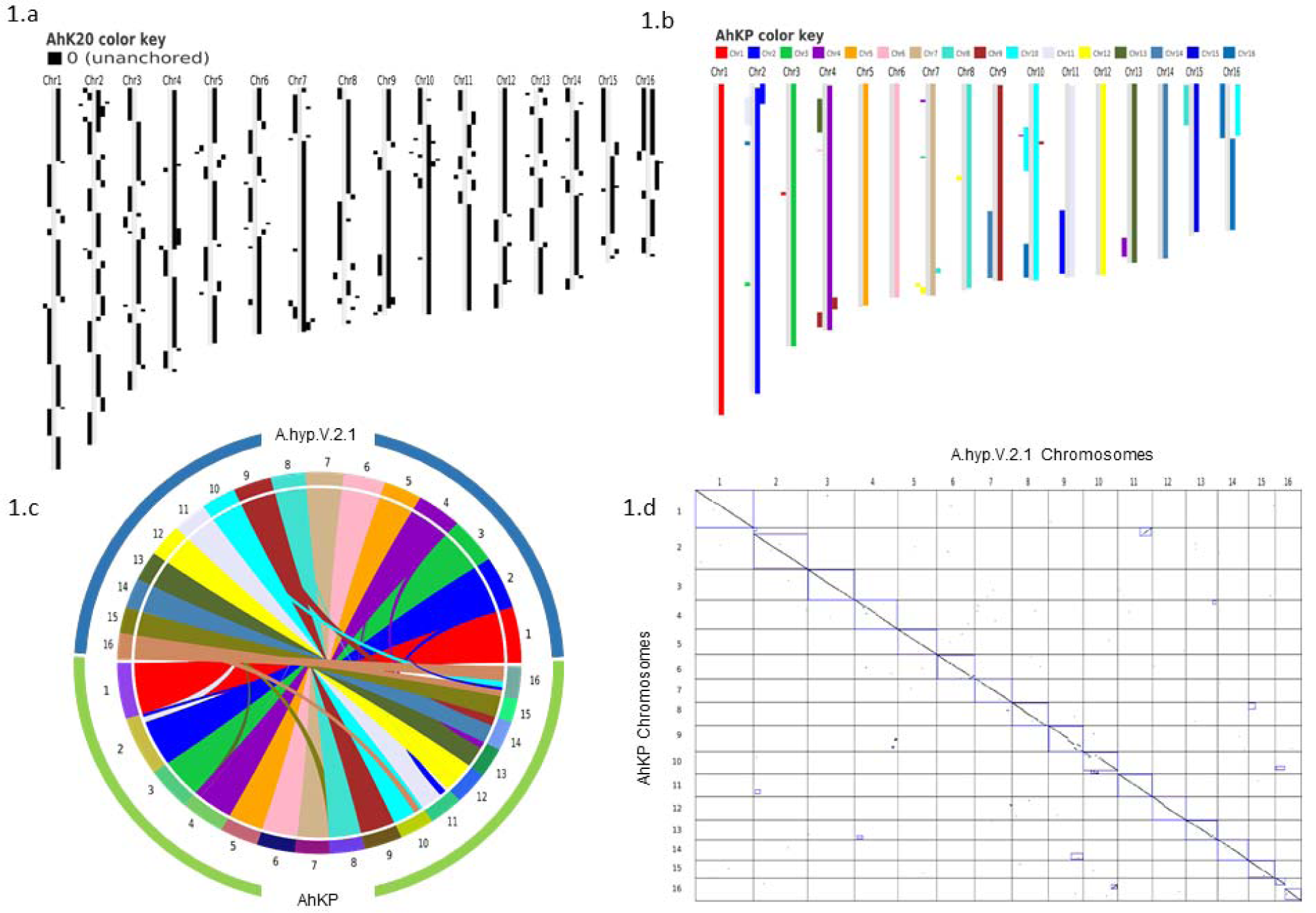
a) Synteny of assemblies with respect to the Plainsman reference A.hyp.V.2.1 against AhK20. b) block plot, c) circular and d) dot plot showing synteny of chromosomes of AhKP assembly on to Plainsman.

### Other landraces and plants

We generated WGS data with coverage of ∼50-150X using the Illumina platform for selected landraces and ornamental varieties. Figure 2 shows the photographs of fully-grown plants sequenced and reported here. These include A.hyp_K_white (Figure 2.a), A.hyp_K_red (Figure 2.b), two ornamental varieties A.cau_ornamental (love-lies-bleeding, Figure 2.c) and A.cru_ornamental (Autumn touch, Figure 2.d) and Suvarna (Figure 2.e). The details of the sequencing are presented in supplementary Table S1. We also downloaded WGS data from NCBI for seven other accessions including A.cau_Bolivia_PI642741, A.cru_Mexico_PI 477913, A.hyp_India_PI481125, A.hyp_Plainsman_PI558499, A.hyp_Nepal_PI619259, A.hyp_Pakistan_PI540446, A.hyp_Mexico_PI511731, and A.hyb_Greece_PI605351. Based on the number of variants called for these and other accessions using both AhKP and A.hyp.V.2.1 as references suggest that all the landraces of *Amaranthus hypochondriacus* sequenced here are different from the A.hyp_Plainsman variety (Supplementary Table S2).

**Figure 2:**
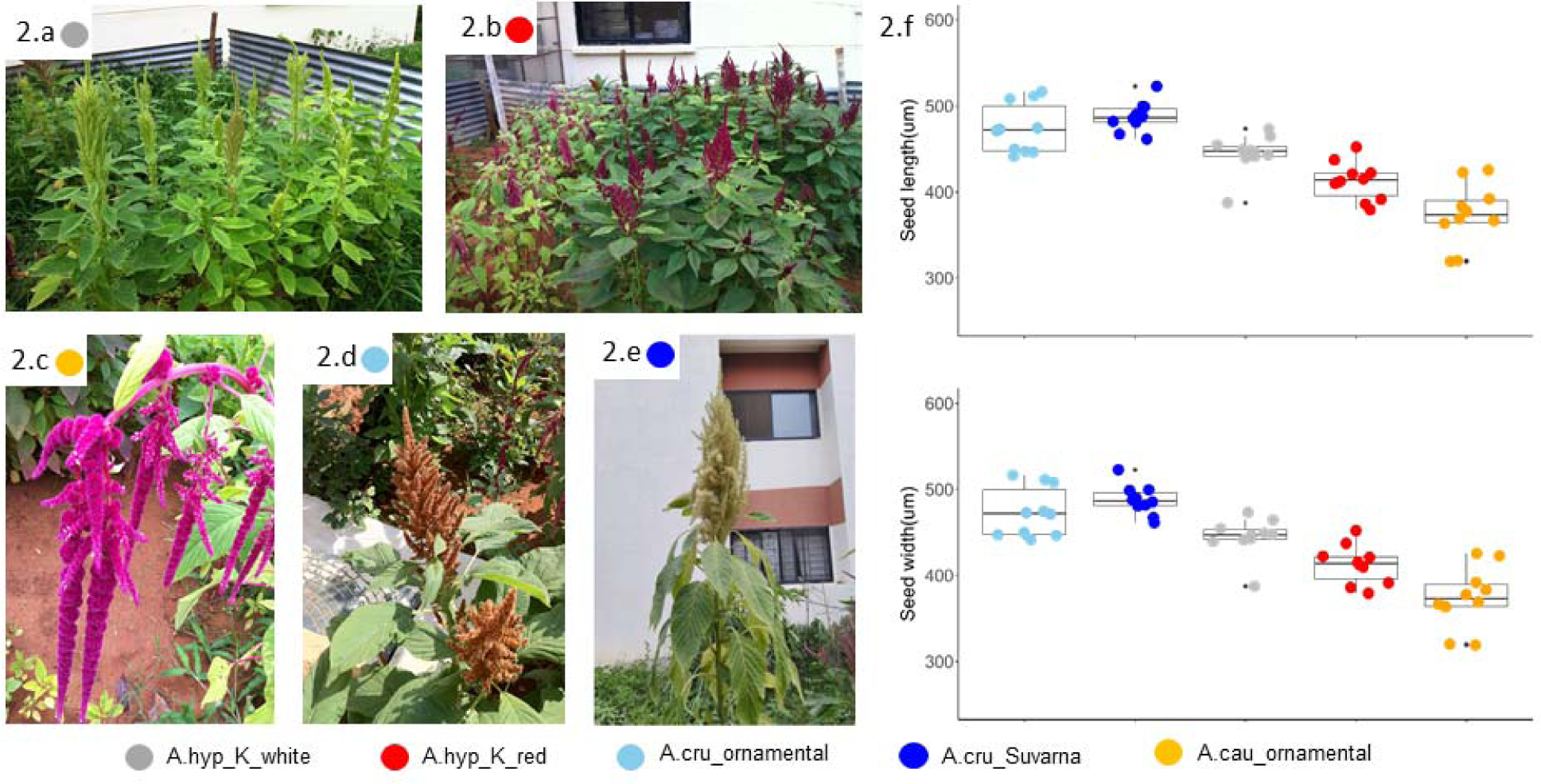
a) Representative images from the institute’s field for the varieties A.hyp_K_white b) A.hyp_K_red c) A.cau_ornamental (love-lies-bleeding) d) A.cru_ornamental (Autumn touch) e) A.cru_Suvarna with white inflorescence grown at the institute campus for taxonomic classification f) color-coded error graph of seed size for each variety.

### Classification of grain amaranths

The variants from WGS data from all the plants in Figure 2 were compared with those from the Plainsman strain and a handful of other accessions from public resources^3^. Figure 3a shows classification using the 27,658 SNPs reported for grain amaranth obtained from Maughan et al.^17^. Of these only 20,548 positions could be found covered in all whole-genome sequencing data studied here.

**Figure 3:**
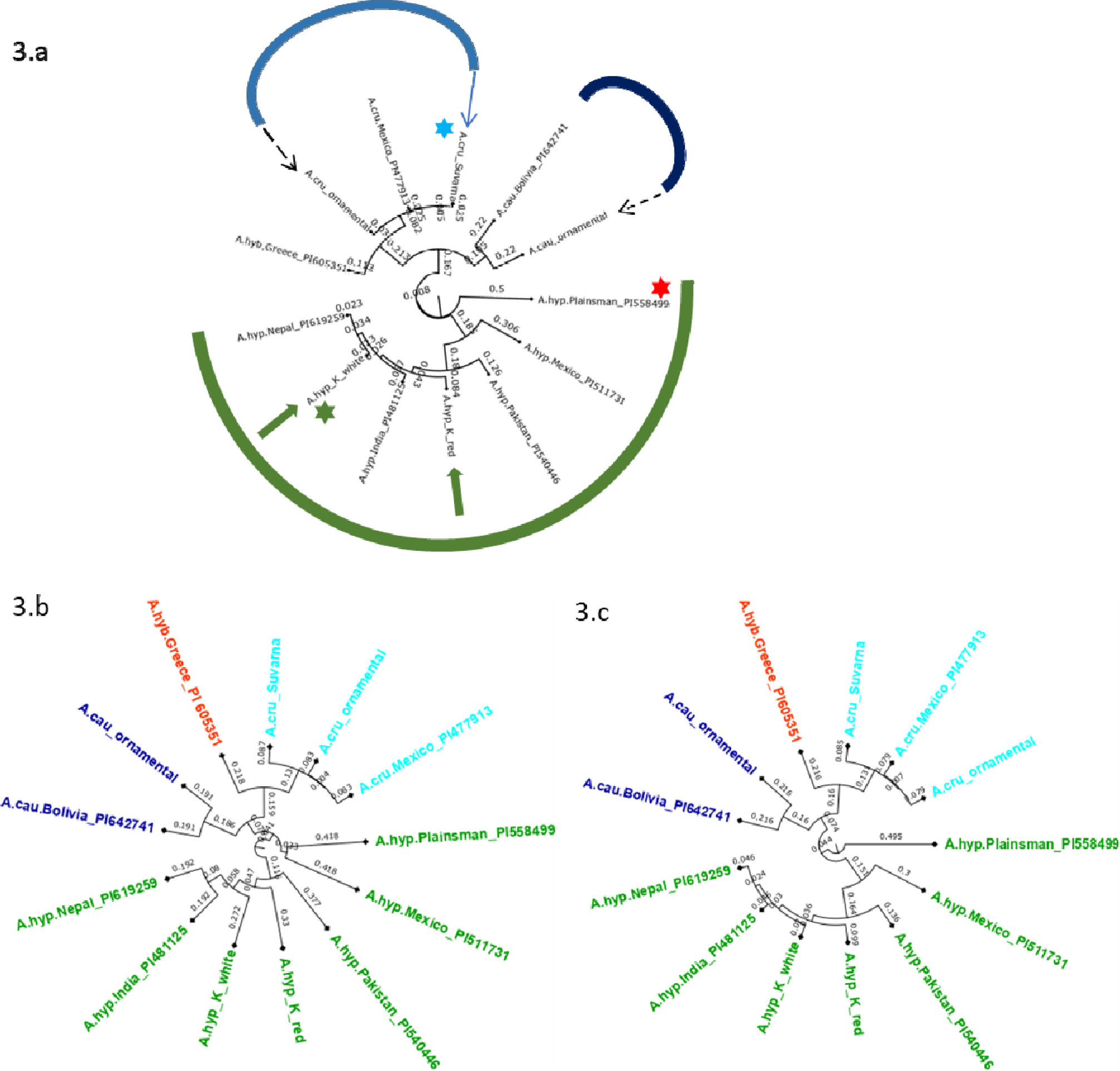
a) Phylogenetic tree using the 20,548 SNPs reported for grain amaranths (green: *A. hypochondriacus*, blue: *A. caudatus* and purple: *A. cruentus*, dashed arrows: ornamental, solid arrows: landraces with green star for A.hyp_K_white and red star for A.hyp_Plainsman variety, Classification using 5,545,132 SNPs from the mapping of short reads onto AhKP as reference, Classification using 6,383,490 SNPs from the mapping of short reads to A.hyp.V2.1 a reference.

All three trees in Figure 3 suggest that A.hyp_Plainsman_PI558499 (red star) is distal to the clade belonging to A.hyp_K_white (green star). The tree generated using the 20,548 SNPs (Figure 3.b) is independent of any reference and hence, can be considered unbiased. On the other hand, for the phylogenetic trees in Figures 3.b and 3.c shows with some bias from the reference used to generate variants also clusters A.hyp_K_white in a distal clade from A.hyp_Plainsman_PI558499. Also, A.hyp_K_white is closest to A.hyp_Nepal_PI619259 and A.hyp_India_PI481125 with A.hyp_K_red relatively distal from A.hyp_K_white. It is interesting to note that A.cau_ornamental clusters close to A.cau_Bolivia_PI642741. Suvarna, an accession/landrace from India, often classified as *hypochondriacus* in the literature, clearly clusters with *A. cruentus and* shows high similarity to the accession A.cru_Mexico_PI477913, also classified as *A. cruentus*. This is also obvious from the stem solidness (Figure 2.e).

In Figure 4.a, an attempt was made to decorate the classification of 94 accessions generated both using GBS data with WGS data for landraces generated here. Since GBS only covers 10% of the genome, there is a need to normalize the variants from WGS data for comparison. One way to do this would be to identify and use the alleles found from GBS data with the respective alleles found from WGS. However, this produced skewed classification because of variation in the depth of sequencing between GBS and WGS while calling variants. We have devised a method to normalize for the same during the variant calling (see methods section).

**Figure 4:**
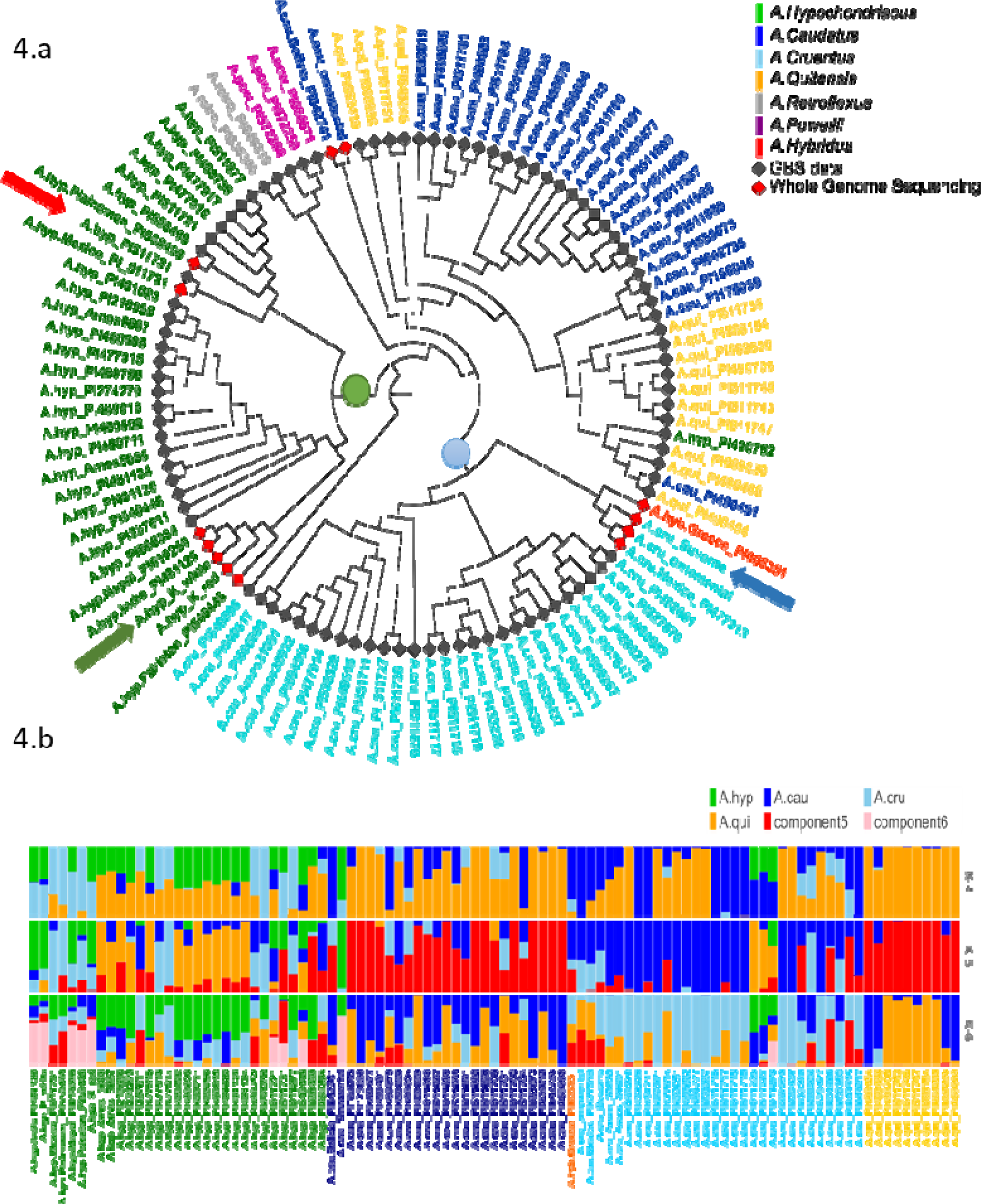
a): Shows a classification of 94 accessions with GBS data and WGS data after normalization of the two sequencing approaches using 271,305 SNPs. b) Genetic admixture analysis of *A.hypochondriacus, A.caudatus, A.cruentus* and *A.quitensis.*

The phylogenetic tree shown in Figure 4.a and generated using AhKP as reference (Figure 4.a) combines variants called for the 94 accessions using both raw genotyping-by-sequencing (GBS) data from public sources^4^ and whole-genome sequencing (WGS) data for listed accessions in supplementary Table S2. The normalization is validated by clustering of both WGS and GBS data from A.hyp_Plainsman_PI558499 and A.hyp_Mexico_PI511731 close to each other (Figure 4.a, red arrow). The tree in Figure 4.a is very similar to that reported using A.hyp.V2.1 as reference^4^. The taxonomy-based classification seems reproducible with green being *A. hypochondriacus*, light blue being *A. cruentus*, dark blue being *A. caudatus* and yellow being *A. quitensis*. However, according to genomics-based classification with 271,305 SNPs, PI490752 is classified as A.*quitensis*, but was originally annotated as *A. hypochondriacus*, and PI649546 is a *A. hypochondriacus* originally annotated as *A. cruentus*. Interestingly, similar to Figure 2, A.hyp_K_white and A.hyp_K_red, both landraces from India cluster closely together along with accessions A.hyp_Nepal_PI619259 and A.hyp_India_PI481125. Suvarna, yet another landrace sequenced and reported here is clearly classified as *A. cruentus*. Figure 2.e shows solid stem characteristics of *A. cruentus* for A.cru_Suvarna. Besides, the seed sizes shown in Figure 2.f also validate classification for Suvarna as *A. cruentus*.

ADMIXTURE analysis shown in Figure 4.b suggests that there is significant gene flow between *A. caudatus* and *A. quitensis*. At K= 4 and 5 there is no resolution between other species. However, at K=6 there is resolution in components for all four species with green for *A. hypochondriacus*, dark blue for *A. cruentus* and major yellow representing components of *A. caudatus* and *A. quitensis*. At K = 6, we also see pink components uniformly present in all the *A. hypochondriacus* from the Indian subcontinent with the exception of accessions A.hyp_Mexico_PI511721(gbs), A.hyp_Mexico_PI511731(gbs). Interestingly, A.hyp_Mexico_PI511721 clusters with A.hyp_Plainsman_PI558499, which does not show any pink components.

### Development of Tissue-specific Gene Expression Atlas of Amaranth

Our lab had sequenced and reported developmental transcriptome of A.hyp_K_white from several tissues^2,5^. Here, the transcriptomes have been mapped to AhKP reference and the expression profiles of the predicted genes have been generated across the developmental stages. The bam files of each sample can be visualized in the respective genome browser (link to the same is available in the data availability section) Also, the expression profiles of all the 12 predicted genes from lysine pathway across developmental stages is provided in Figure 5.a along with the corresponding sizes compared to Arabidopsis Figure 5.b. Also, the browser can be queried using the accessions of Arabidopsis to visualize the expression profile of the corresponding orthologs on AhKP.

**Figure 5:**
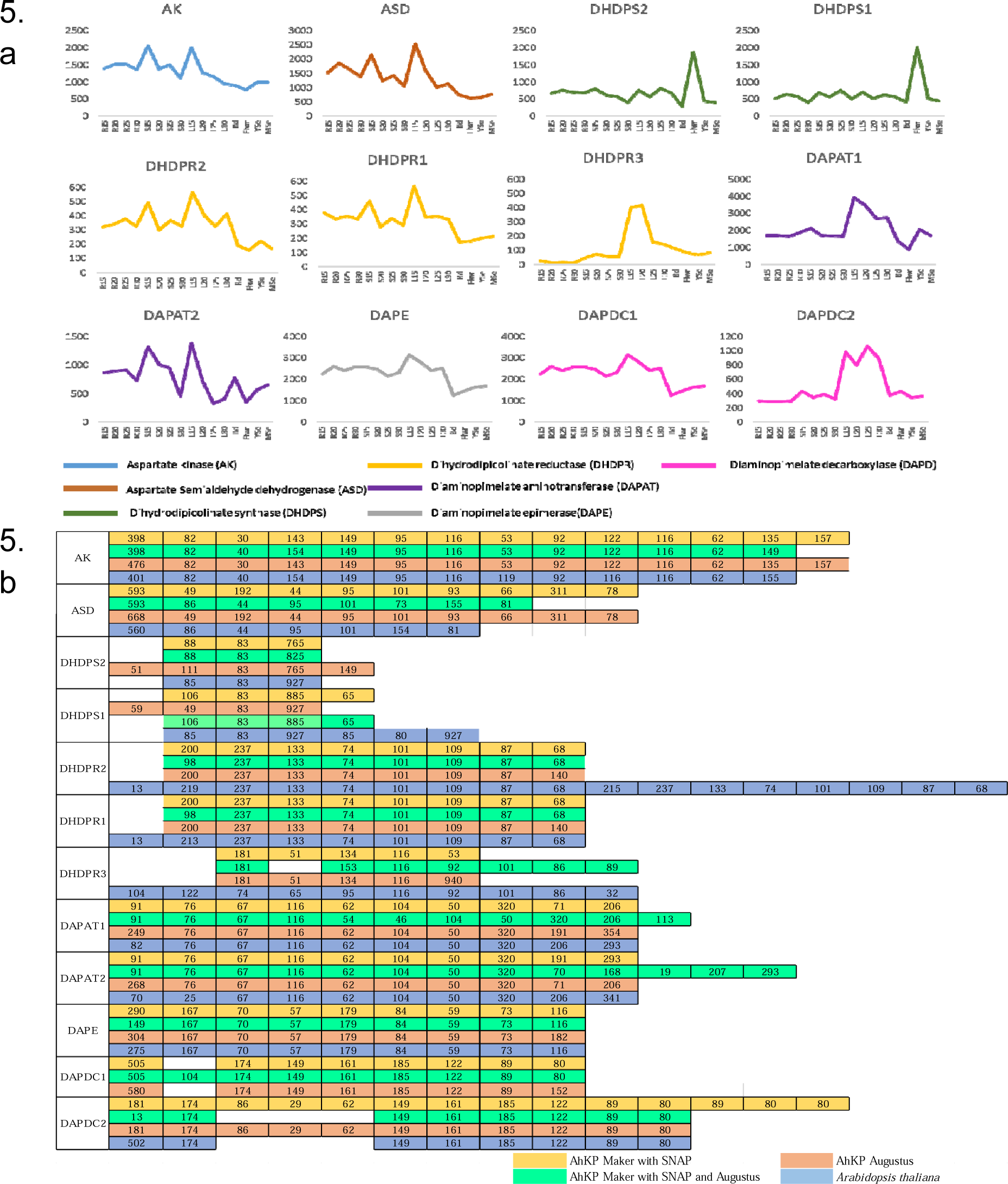
a) Gene expression profile of 12 genes involved in lysine biosynthesis pathway across different developmental stages (15, 20, 25, 30 days) of different tissues (Rt-root, St-stem, L-leaf, Bud, Flwr-Flower, YSe-young seed, MSe-mature seed). b) Comparison of CDS sizes (nucleotides) of lysine biosynthesis pathway genes predicted in AhKP with Arabidopsis.

**Figure 6:**
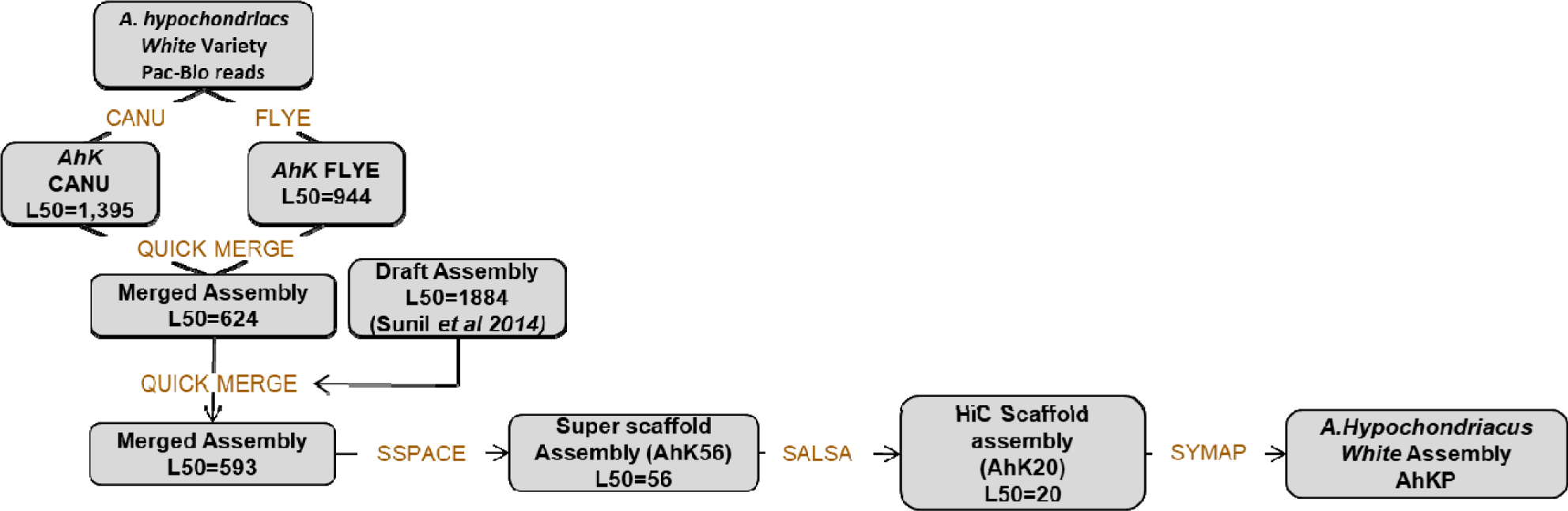
Workflow used in the assembly of AhKP.

## DISCUSSION

Here, a chromosome level assembly (AhKP) of a landrace, A.hyp_K_white, under contiguous cultivation in India for over several centuries is reported for which, a draft genome was reported by our group in 2014^2^. The assembly reported here is obtained using relatively low coverage of long reads from PacBio RSII technologies in conjunction with a high-quality reference for another distal strain of the same species A.hyp_Plainsman_PI558499. We used multiple assembly tools adapted/developed recently to handle error-prone long reads and merged these assemblies with the contigs from our previously reported draft assembly. The assembly statistics of the initial assembly was sufficient for reference-based scaffolding using both the simulated mate-pairs from the reference genome and raw HiC data for Plainsman from public resources^3^. RepeatMasker analysis classified 50.5% (196421031 bp) of the AhKP genome as repetitive sequences. Annotation using the MAKER annotation pipeline predicted 18,858 gene models which has been validated for the 12 genes from lysine biosynthesis pathway by comparing it to Arabidopsis gene model as shown in Figure 5.

Also, we measure the extent of the diversity of A.hyp_K_white and a few other landraces from India with respect to other known accessions. Whole-genome sequencing from a few landraces and ornamental varieties generated in-house and that for several other accessions from public sources are clustered using 27,658 SNPs reported for grain amaranths (Figure 3.a). Figure 3 shows classification using both the 20,548 out of 27,658 reported SNPs covered in all samples (Figure 3.a) and ∼6 million variants called from mapping to AhKP and A.hyp.V2.1 reference respectively. In that, the genome of the landrace AhKP, reported here, is closest to accession A.hyp_Nepal_PI619259 and to A.hyp_India_PI481125. However, A.hyp_Plainsman_PI599488 clusters in a distal clade with A.hyp_Mexico_PI511731 in both Figure 3 and Figure 4. Also, the two landraces under hypochondriacus, A.hyp_K_white and A.hyp_K_white, cluster apart. This validates our observation that the seeds of these two varieties faithfully produce inflorescence with the same color as the parent plant. Besides, a C0t analysis shown in supplementary Figure S2 suggests distinct dissociation time for simple repeat between these two accessions.

The successful integration of WGS and GBS data attempted here, is apparent from the clustering of variants from WGS and GBS data for the same accession together as marked with red arrow in Figure 4a. Figure 4 validates taxonomy-based classification (colour-coded accessions) of the majority of the accessions and landraces. However, a few accessions are now reclassified. The accession PI490752 originally classified as *A. hypochondriacus* now classifies under *A. quintensis. A. hybridus* with accession of PI605351 clusters in the same clade as Suvarna with another accessions (PI477913) from *A. cruentus*. All the accessions from *A.quintesis* and *A.caudatus* clusters together in a single clade with two branches of *A. quintesis* enclosing *A. caudatus,* which is also reported using only GBS data using A.hyp.V.2.1 as reference^4^. This suggests that *A. caudatus* is a major clade under *A. quintesis*. ADMIXTURE analysis shown in Figure 4.b, also suggests that there is significant gene flow between *A. caudatus* and *A. quitensis*. At K= 4, a significant component of *A. quintesis* is found in all four species. However, at K= 5 and 6 components of *A. quintesis* is profound only in *A. caudatus*. At K= 6 other unique components within *A. hypochondriacus* gets resolved. For example, there is a component (Figure 4.b, pink) only present in all *A. hypochondriacus* from India, which is missing in Plainsman.

Suvarna (R 104-1 −1), a pureline released by University of Agricultural Sciences (UAS), Bangalore-1992 from the material ‘Rodale Plus’ received from Rodale Institute^18^ has previously been classified as *A. hypochondriacus* in an article titled “Evaluation of Genetic Diversity in Grain Amaranth (*Amaranthus hypochondriacus*) at Molecular Level using ISSR Markers” using ISSR markers and other classification techniques^19^. Here, Suvarna is undoubtedly classified as *A. cruentus* based on 20,548 reported SNPs and roughly 6 million variants covered in WGS data (Figure 3). Also, morphological features like stem solidness (Figure 2.e) and seed size (Figure 2.f) supports this classification. The total height for Suvarna at maturity reaches 8-9 feet compared to ∼6 feet for both red and white varieties of *A. hypochondriacus* reported here. In Figure 2, the height and stem solidness of Suvarna are very similar to *A. cruentus* but the inflorescence is similar to *A. hypochondriacus*, which may be the reason for the misclassification of Suvarna as *A. hypochondriacus* in the literature. We hypothesize that the only component showing light-blue that is common between Suvarna and A.hyp_K_white in the ADMIXTURE with K = 5 and 6 (Figure 3b) holds the genotype responsible for inflorescence within this haplo-block.

We believe that this is the first demonstration of generating a cost-effective de novo assembly for a landrace utilizing low coverage PacBio reads in conjunction with the genome and HiC data from another strain. Since this landrace is more closely similar to all other landraces and accessions for *hypochondriacus* from India and South Asia (supplementary Table S1), AhKP offers a better reference for the improvement of grain amaranth crops in South Asia. The landrace A.hyp_K_white is currently being used to identify mutations in targeted loci for a given desirable phenotype from a germplasm collection using eco-TILLING and to discover novel mutations that result in desirable traits using TILLING-based approaches.

## MATERIALS and METHODS

### Samples

Seeds of A.hyp_K_white, A.hyp_K_red were obtained from local market in Karnataka, India, A.cru_ornamental, A.cau_ornamental from Park seeds and A.cru_Suvarna from Gandhi Krishi Vigyana Kendra (GKVK), Bengaluru, Karnataka, India.

### Source of data used in this work

Plainsman reference: Phytozome (http://phytozome.jgi.doe.gov/) *Amaranthus hypochondriacus*

genome V.2.1(A.hyp.V.2.1)

GBS: Blair et al Front Plant Sci. 2017

WGS: https://www.ncbi.nlm.nih.gov/sra?term=SRP061623

### Isolation of Genomic DNA

Amaranth A.hyp_K_white, A.hyp_K_red, A.cru_ornamental, A.cau_ornamental and A.cru_Suvarna variety were grown at IBAB (Figure 2). Genomic DNA was extracted from fresh leaves using the DNeasy Mini Plant DNA Extraction kit (Qiagen) following the manufacturer’s protocol and quantified using fluorometry (Qubit 2.0, Invitrogen).

### Library preparation and sequencing

Whole Genome libraries were prepared using the TruSeq DNA Sample Preparation Kit (Illumina) by following the manufacturer’s low throughput protocol. One microgram and 10 µg of the DNA were used for the preparation of Paired-End (PE) and Mate-Pair (MP) libraries, respectively. DNA was sheared using Adaptive Focused Acoustic technology (Covaris, Inc.) to generate fragments of desired insert size. The average insert size was around 200 bp for PE libraries and 1.75, 3, 5, and 10 kb for four MP libraries.

Briefly, for PE libraries, the fragmented DNA was end-repaired, 3′-adenylated, ligated with Illumina adapters, and PCR enriched with Illumina sequencing indexes. For MP libraries, the fragmented DNA was end-repaired, followed by end labeling using the biotin-dNTP mix, size selected and later, circularized using circularization ligase. The circular DNA was sheared again as explained earlier, and the biotinylated fragments were purified using streptavidin beads (Dynabeads(tm) M-280 Streptavidin, Invitrogen), the fragments were end-repaired, 3′-adenylated and ligated with Illumina adapters. Further, the biotinylated, adapter-ligated immobilized DNA were enriched by PCR. The size selection for all the libraries were done using solid-phase reversible immobilization (SPRI) beads (Agencourt AMPure XP Beads) from Beckman Coulter. The quality, quantity, and size distribution of the libraries were evaluated using Qubit (Invitrogen) and TapeStation (Agilent)^2^. The clusters were generated in cBot and paired-end sequenced on Illumina HiSeq 2500 platform.

Whole-genome PacBio sequencing was done by Molecular Biology and Genomic Core, Washington University using P5/C3 chemistry on the Pacific Biosciences RSII platform. This platform is a single-molecule, real-time (SMRT) sequencing machine that uses a sequencing-by-synthesis method to generate good quality very long reads.

### Assembling the raw data

The raw PacBio data was assembled using Canu^13^ and Flye^14^ independently. The two assemblies obtained were then merged together using Quickmerge^15^. This was further improved by merging the Illumina assembly from the draft genome reported elsewhere and polished using the Illumina reads. The scaffolds from this step are further improved with simulated mate pairs using wgsim^20^ from Plainsman with SSPACE. At this stage, the scaffolds were long enough to allow the use of HiC data generated for Plainsman to obtain high-resolution assembly (AhK20). Further, we generated synteny of the AhK20 against A.hyp.V.2.1 using Symap^21^ based on which AhK20 was improved to the final AhKP assembly. The flowchart below (Figure 6) shows the pipeline used to obtain the final assembly.

### SNP analysis and construction of phylogenetic tree for whole genome samples

The Illumina data of all the plants with accessions listed in supplementary Table S2 were downloaded from NCBI SRA (SRP061623). The public and in house generated data were mapped to A.hyp.V.2.1 and AhKP reference using bowtie2^22^. From the mapped reads, variants were called using samtools (v1.9) mpileup^23^ and bcftools (v1.9)^23^. The variants were filtered using bcftools^24^ with the criteria of QUAL (quality) greater than 10 and DP (read depth) greater than 3 and INDELs were also removed. The files were then merged, and the genotype matrix was created using a custom script. Further, the regions covered in sequencing were identified using bedtools genomecov^24^ from bam files, and the regions, which were not covered in sequencing in any of the samples, were removed from the genotype matrix. The phylogenetic tree was constructed from the genotype matrix using the clustering algorithm hclust and SNPRelate under R and Bioconductor package^25^.

For the 27,658 SNP positions 150 base pair sequences were downloaded from public sources and coordinates for all 27,658 positions on A.hyp.V.2.1 were extracted by BLAST alignment. A separate VCF file was made for all the 13 datasets as listed in supplementary Table S2 with the respective alleles at these positions. Only 20,548 SNPs were commonly covered in all 13 WGS datasets and were used during classification. The resultant VCF files were merged and based on the presence or absence of SNPs, a binary matrix was constructed from which a phylogenetic tree was obtained as mentioned above.

### Classification using GBS and WGS data

GBS raw data of 95 accessions were downloaded from Blair et al ^4^ of which *A. palmeri* was excluded from the analysis because of the reported high level of missing data. The reads were demultiplexed using GBSX^26^ using the provided barcode sequences. Post demultiplexing, the reads were mapped to AhKP using bowtie2^22^ and SNP calling was done using the method described in the above section.

To combine WGS data and GBS data, we created GBS like data from whole-genome reads. For this, the regions covered in GBS data were extracted using bedtools genomecov^24^ for all the accessions, and the regions covered were merged to get a maximum possible region covered in GBS sequencing for all the accessions. These regions were used to restrict the variant calling from whole-genome data to only the regions covered in GBS. Also, the read depth considered during variant calling was restricted to 10 to match the depth of GBS data^24^. The SNPs were merged and used for phylogenetic classification.

### Admixture Analysis

Population genetic diversity was analyzed for four Amaranth species (*A. hypochondriacus, A. caudatus, A. cruentus and A. quitensis).* Only 97 out of 107 samples from both GBS and WGS data were filtered based on their good clustering and bigger sample size. The merged SNP file was processed using PLINK^27^ and Admixture (v1.3) was used to analyze the population structure^28^.

### Genome annotation and repeat analysis

Repeat elements for the Plainsman and the A.hyp_K_white variety of *A. hypochondriacus* assemblies were predicted using RepeatModeler version 2.0.1^29^ along with LTR discovery. The two predicted libraries of repeat elements were merged together and repeat masking was done using RepeatMasker version 4.1.0^30^.

Annotation of AhKP was done using multiple approaches i) Augustus^31^ (v3.2.3) prediction using *Arabidopsis* as model and ii) MAKER^32^ genome annotation pipeline with(with/without Augustus) default parameters, was used for AhKP annotation. Maker pipeline includes de novo assembled amaranth transcriptome with 125581 scaffolds, repeat elements predicted by RepeatModeler and *Arabidopsis* proteome (TAIR10)^33^. SNAP^34^ and Augustus were also used to predict gene models and used in the subsequent rounds of MAKER^32^.

Genes involved in lysine biosynthesis pathway were identified by BLASTP^35^ analysis using *Arabidopsis* proteins.

### Transcriptome analysis

Raw transcriptomic reads from 16 developmental stages were mapped to AhKP reference using bowtie2^22^. The mapped files were processed using samtools^23^ and raw read count was counted for all predicted genes using bedtools multibamcov^24^. Further DESeq2^36^ was used to get normalized read counts.

### Genome browser and database

The Amaranth database is running on EC2 instances of Amazon cloud service (AWS). The database is built using HTML5, bootstrap and Javascript. The database consists of a landing page, genome browser and BLAST tool. This database is made from a framework provided by Meghagen LLC. Jbrowse^37^ is javascript and html based genome browser provides the solution for visualization of various kinds of genomic data such as FASTA, BAM, GFF, VCF and bigwig etc. Data for downloading and JBrowse is stored on the cloud and made available for research purposes. The menus on the database page will redirect you to the download as well as tool page. Users can access the Jbrowse by clicking on the Genome browser button or using the tools menu. Users can access the database from the link given in the data availability section. The database is also integrated with graphical visualization for gene expression data of 16 developmental stages with query search options.

## ACKNOWLEDGEMENTS

The authors wish to acknowledge GKVK for providing us with seeds for Suvarna and to Dr. Xingbo Wu of Dr. Blair’s lab for providing us with raw GBS data from 94 accession. The authors wish to recognize lab infrastructure support from DST, computing infrastructure by GoK and DBT for support to Saptarathi Deb via JRF under the project BT/PR23613/BPA/118/354/2017 titled “Non-transgenic crop improvement of grain amaranths (*A. hypochondriacus*) for determinate growth, enhanced seed yield and oil by establishment of TILLING by sequencing platform".

## DATA AVAILABILITY

http://52.4.112.252/ (**Amaranth Repository / Database and genome browser)**

## AUTHOR CONTRIBUTIONS

SPD: Classification, characterization and writing of the manuscript; SJ: Assembly of AhKP; SR: Library preparation and aiding writing of manuscript; RRK: Assembly and analysis of other landraces; SW: Development of genome browser; NH: Transcriptome analysis; SD: DNA isolation and repeat analysis; MS: PacBio data, developmental transcriptome and taxonomic classification of landraces; ER: For validating transcripts; BC: For overseeing the experimental component of the project; PGS: For guidance throughout the project; SS: For overseeing the project and writing of the manuscript

