## Supplementary Material for "Classification of grain amaranths using chromosome-level genome assembly of ramdana, *A. hypochondriacus*"

**SUPPLEMENTARY DATA**

Figure S1 - *A. hypochondriacus* Plainsman PI558499

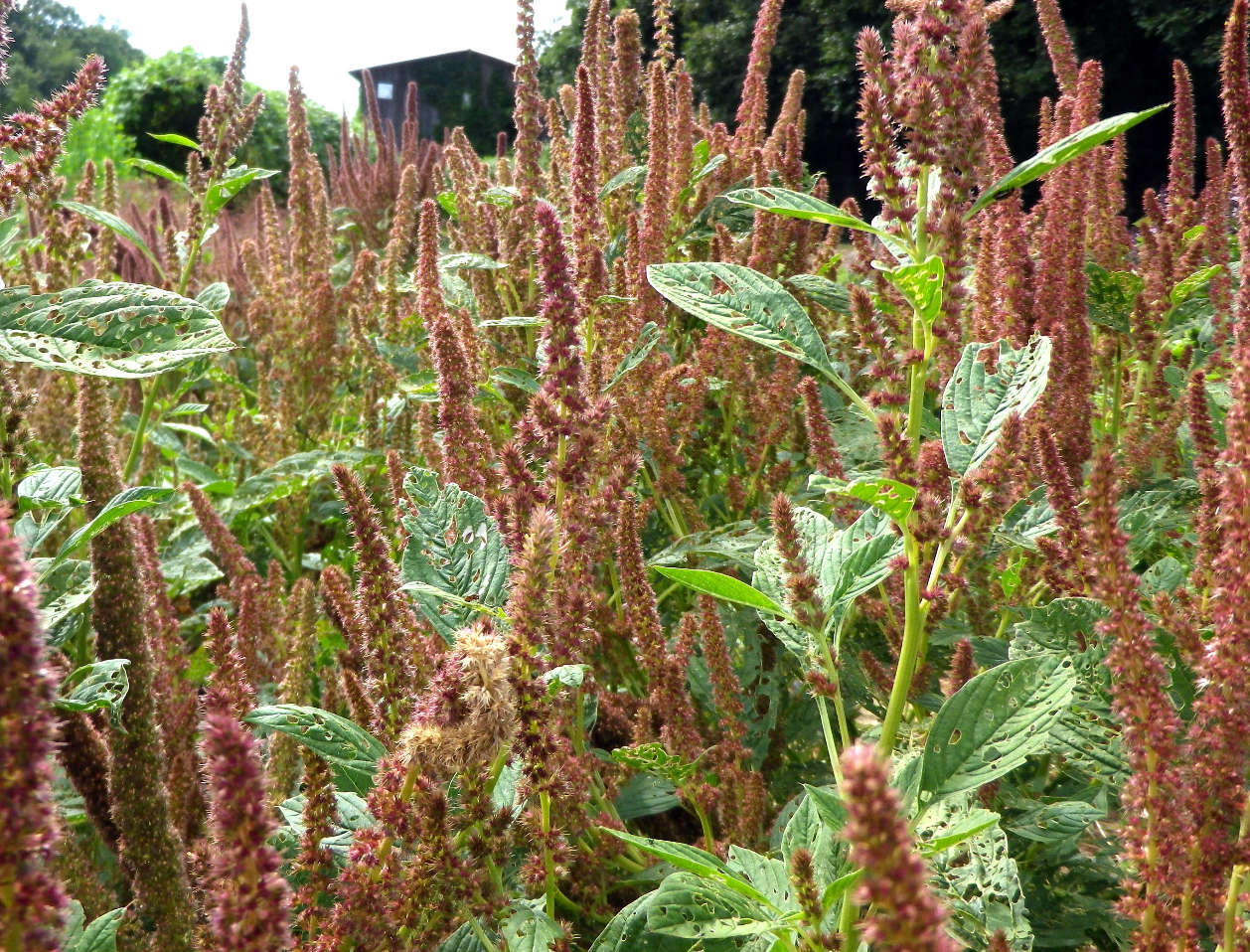

Figure S2 - C_0_t analysis of A.hyp_K_white and A.hyp_K_red

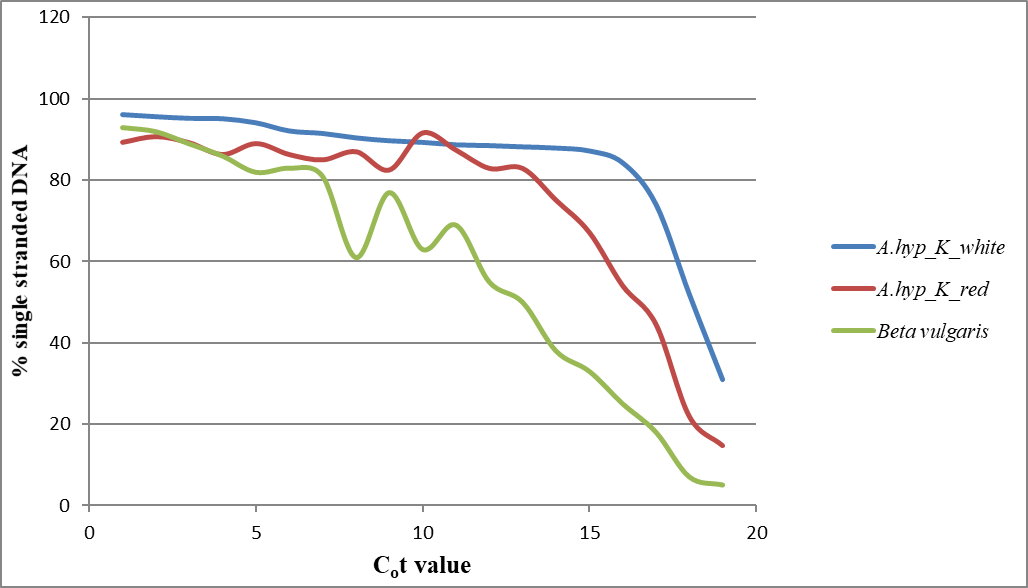

TABLE S1- Details of in-house sequenced plants

|  | **Total no of reads** | **Coverage** |
| --- | --- | --- |
| **A.hyp_K_white** | 469749648 | 87.3357055 |
| **A.cau_ornamental** | 266702040 | 53.1278964 |
| **A.cru_ornamental** | 353363298 | 69.2869212 |
| **A.hyp_K_red** | 447933760 | 111.03960 |
| **A.cru_Suvarna** | 501687774 | 147.555228 |

TABLE S2- SNP count of all whole genome samples on both references

| **Name** | **Alias** | **On A.hyp.V.2.1** | **On AhKP** |
| --- | --- | --- | --- |
| *A. caudatus* (Bolivia) PI 642741 | A.cau_PI642741 | 2846014 | 2424857 |
| *A. cruentus* (Mexico) PI 477913 | A.cru_PI477913 | 3263942 | 2773417 |
| *A. hypochondriacus* (India) PI481125 | A.hyp_PI481125 | 886235 | 117514 |
| *A. hypochondriacus* K white (India) | A.hyp_K_white | 901042 | 110024 |
| *A. hypochondriacus* Plainsman PI558499 | A.hyp_PI558499 | 78926 | 765147 |
| *A. hypochondriacus* (Nepal) PI619259 | A.hyp_PI619259 | 881424 | 112943 |
| *A. hypochondriacus* (Pakistan)PI540446 | A.hyp_PI540446 | 937366 | 196203 |
| *A. hypochondriacus* K red (India) | A.hyp_K_red | 1006061 | 193013 |
| *A. hypochondriacus* (Mexico) PI511731 | A.hyp_PI511731 | 1248769 | 738991 |
| Suvarna (India) | A.cru_Suvarna | 3267631 | 2796817 |
| *A. hybridus* (Greece) PI605351 | A.hyb_PI605351 | 3280993 | 2783846 |
| *A. caudatus* (Love-Lies-Bleeding) | A.cau_ornamental | 2954432 | 2138832 |
| *A. cruentus* (Autumn Touch) | A.cru_ornamental | 3428292 | 2920539 |
